# Role of DNA Methylation in Persister Formation in Uropathogenic *E. coli*

**DOI:** 10.1101/2020.01.07.897686

**Authors:** Yuanyuan Xu, Shuang Liu, Ying Zhang, Wenhong Zhang

**Affiliations:** Department of Infectious Diseases, Huashan Hospital of Fudan University, Shanghai, China; Department of Molecular Microbiology and Immunology, Bloomberg School of Public Health, Johns Hopkins University, Baltimore, MD, USA

**Keywords:** Persister, DNA adenine methylation, Methylome, Transcriptome, Urinary tract infection

## Abstract

Uropathogenic *Escherichia coli* (UPEC) persister bacteria play crucial roles in clinical treatment failure and relapse. DNA methylation is known to regulate gene expression in bacteria, but its role in persister formation has not been investigated. Here, we created adenine methylation deletion mutant (*Δdam*) and cytosine methylation mutant (*Δdcm*) from UPEC strain UTI89 and found that the *Δdam* mutant but not *Δdcm* mutant had significant defect in persister formation during exposure to various antibiotics (gentamicin, fluroquinolones and cephalosporin) and stresses (acid pH and hyperosmosis), and that complementation of the *dam* mutant restored its persister defect phenotype. PacBio sequencing of epigenetic genomewide methylation signature and RNA sequencing of the *Δdam* mutant were performed to define, for the first time, the role of adenine methylation in persister formation. Methylome data analysis showed that 99.73% of m^6^A modifications on GATC were demethylated in the *Δdam* mutant, and demethylation nucleotide site related genes suggested an overwhelming effect on transcription and metabolic processes. Transcriptome analysis of the *Δdam* mutant in comparison to wild type showed that flagella biosynthesis, galactitol transport/utilization, and signaling related genes were upregulated while pilus, fimbriae, virulence, glycerol, nitrogen metabolism pathways and transcriptional regulators were downregulated. The comparative COG analysis of methylome and transcriptome enriched pathways identified translation, ribosomal structure and biogenesis, and cell motility were upregulated, whereas DNA repair, secondary metabolite biosynthesis and diverse transport systems, some of which are known to be involved in persister formation, were downregulated in the *Δdam* mutant. These findings provide new insights about the molecular basis of how DNA adenine methylation may be involved in persister formation and offer novel therapeutic targets for combating persister bacteria.

## Introduction

Urinary tract infections (UTIs) are among the most common infectious diseases worldwide. UTIs often result in recurrent UTI (RUTI) that last weeks or months in patients, and almost half of all women will experience a UTI in their lifetime ^1^. The most common causative agent of UTIs is uropathogenic *Escherichia coli* (UPEC) ^2^. The long-term exposure or frequent retreatments can result in development of antibiotic resistance ^3^. Type 1 pilus mediated UPEC entry into bladder epithelial cells and invasive adhesins exist in virtually all UPEC isolates ^4^. Once internalized in bladder epithelial cells, the bacteria can persist for a long time in a quiescent state, and this process enables the persistent bacteria, also called intracellular bacterial community (IBC) to escape host cell elimination and resurge from the reservoirs. Some studies estimated that 50-78% recurrent strains are genetically identical to the original strains, indicate the persistence of the original strain ^5 6 7^. The RUTI cases were defined as “persistence” of the same strain as the initial strain caused the infection ^8^.

“Persisters”, which were discovered more than 70 years ago ^9^, are a subpopulation of dormant and metabolically quiescent bacteria that are tolerant to antibiotics and stresses. UPEC bacteria can transform into persisters, which can exhibit tolerance to not only antibiotics but also host cell defenses ^10^. Persistence or tolerance is due to epigenetic changes and differs from genetic antibiotic resistance. Antibiotic tolerance can facilitate subsequent development of antibiotic resistance ^11 12^. In addition, the presence of persister bacteria is an important cause of relapse and complicated UTIs clinically ^10^. Typically, IBCs and biofilm structures which are embedded in self-secreted extracellular matrix (ECM) containing dormant persister cells can act as reservoirs for persistent infections and relapse once antibiotic stress is removed ^13^ and can pose significant challenges for clinical treatment ^10^.

Persisters are phenotypic variants that are formed through epigenetic changes that involve multiple redundant pathways to survive lethal antibiotic stress. These include toxin-antitoxin (TA) modules, stringent response (ppGpp) ^14, 15^, DNA repair, metabolism ^16^, energy production, protein degradation pathway (trans-translation), signaling pathways, antioxidant system, and efflux and transporters ^17 18^. To better understand persister cells development and functioning, classical genetic processes is not sufficient, the epigenetic controlling gene expression is urgently required. The primary epigenetic-modification system is made of a restriction endonuclease and DNA adenine or cytosine methyltransferase. In *E. coli*, DNA cytosine methylase (Dcm, encoded by *dcm*) is not known to be involved in gene regulatory control whereas DNA adenine methylase (Dam, encoded by *dam*) plays an essential role in regulating epigenetic circuits as the orphan methylase ^19^. Dam methylates adenine of 5’-GATC-3’ motif in newly synthesized DNA to m^6^A (N^6^-DNA adenine methylation) ^20^, which can regulate various important biological processes, including transcriptional control, mismatch repair, chromosome replication and host-pathogen interactions ^19, 21^. In *E. coli* and its relatives, lack of Dam methylation causes pleiotropic defects, indicating that the multiple DNA-protein interactions are under GATC methylation control ^22 23^. DNA methylation has been found to be required in biofilm production as absence of Dam in *S. enteritidis* affected the ability to form biofilm on polystyrene surfaces ^24 25 26^. In addition, it has been reported that a tautomerizing demethylation could disrupt the biofilm formation in anoxic conditions in *P. aeruginosa* ^27^. Furthermore, a biphasic epigenetic switch-N(6)-adenine DNA-methyltransferase (ModA) in *Haemophilus influenzae* was found to control biofilm formation, where Dam ON formed more robust biofilms than Dam OFF ^28^. Persister cells are known to contribute to the antibiotic tolerance observed in biofilms ^29^. However, the role of *de novo* DNA methylation in persister formation is unknown. In this study, we hypothesized that epigenetic modification by DNA methylation may affect the persister phenotype generation.

To address the role of DNA methylation in persister formation, we constructed *Δdam* mutant and *Δdcm* mutant in uropathogenic *E. coli* strain UTI89 and found that, compared with the parent strain, the *Δdam* mutant but not had significant defect in persister formation. We also performed high-resolution DNA methylation analysis of the entire genome (methylome) as well as transcriptome analysis of the *Δdam* mutant in comparison with the parent strain UTI89. Methylome analysis showed numerous genomic positions were demethylated in the *Δdam* mutant. Genome-wide clusters of orthologous gene (COG) distribution analysis indicated predicted genes are involved in transcription and metabolic pathways. Transcriptome analysis revealed that galactose/propanoate metabolism and flagellar assembly were preferentially overexpressed, whereas the transcription regulator, sugar phosphotransferase systems (PTS) and secretion-related pathways were down-regulated in the *Δdam* mutant. The COG comparison revealed key pathways enriched in transcriptional control, cell motility, DNA repair process and secondary metabolite transport to be involved in Dam-mediated persister formation. Together, our findings define an important role for DNA methylation in persister formation in *E. coli.*

## Material and Methods

### Bacterial strains and growth conditions

The bacterial strains used in this study were UPEC strain UTI89 and its derivatives and were routinely cultured in Luria-Bertani (LB) broth at 37□ with shaking (200 rpm) or on LB agar after incubation overnight. Bacterial stock stored at −80□ was transferred into fresh LB medium and grown overnight before use. Ampicillin (Amp^R^), chloramphenicol (Cm^R^) and tetracycline-resistant (Tet^R^) transformants were selected on LB agar plates containing the respective antibiotic at the concentration of 100, 25 and 10 μg/ml. Strains, plasmids and primers used in this study are listed in Supplementary Table 1.

### Construction of *E. coli* UTI89 *dam* knockout mutant and its complemented strains

Disruption of *dam* in the *E. coli* chromosome was achieved by λ-red recombination system as previously described ^30 31^. Primers designed for the purpose are listed in Supplementary Table 1. Briefly, knockout-DNA fragments were generated by using primers KO-dam-F/R with 50nt extensions that are homologous to regions adjacent to the *dam* gene from UTI89 and template chloramphenicol resistant cassette (cat) from plasmid pKD3, which was flanked by FRT (flippase [FLP] recognition target) sites. The above PCR product was electroporated into the competent UTI89 (*dam+*) cells contained pKD46 plasmid carrying λ-red recombinase. The transformants on LB (Cm^R^) plates were selected and verified by primers V-dam-F/R. For complementation experiments, the *dam* gene operon, including its natural promoter and terminal regions, was cloned into plasmid pBR322. The new constructs along with the empty vector control pBR322 were transformed into the *dam* mutant competent cells by electroporation ^32^. The complemented strain and plasmid insertions were confirmed by PCR and DNA sequencing. Meanwhile, we also constructed the *dcm* knockout mutant in the UTI89 to investigate correlation between persister formation under various antibiotic stresses.

### Persister assay

Persister assay was performed by determining the bacterial survival as colony-forming units (CFUs) per ml after antibiotic exposure as previously described ^33^. Stationary phase cultures (10h) were exposed to various antibiotics including 40 μg/ml gentamicin, 10 μg/ml levofloxacin, 10 μg/ml ciprofloxacin, 96 μg/ml cefalexin 8 μg/ml norfloxacin and 200 μg/ml ampicillin for various times. The initial cell number was determined by sampling 10 μl and serially dilution in phosphate-buffered saline (PBS), washing before plating on LB agar. The CFU counts were measured after overnight incubation at 37□.

### Susceptibility to various stresses

For heat shock, bacterial cells were placed in a water bath at 52□ for 1.5h. For acid stress (pH 3.0) and hyperosmosis (NaCl, 4M), bacterial cultures were washed twice with acid or hyperosmotic LB medium and resuspended in the same volume of corresponding medium. Aliquots of exponential and stationary phase cultures were treated with various stresses and were incubated at 37□ at different time points and washed before plating on LB plates in the absence of antibiotics to measure CFU count.

### PacBio SMRT sequencing and DNA methylation analysis

Bacterial DNA methylation analysis was performed by SMRT (Single molecule, real-time, PacBio) and high throughput DNA *de novo* sequencing. Detection of base modification and methylated motif was conducted by employing the SMRT analysis portal following genome assembly. Briefly, genomic DNA was extracted from stationary phase culture of *E. coli* UTI89 and *dam* mutant, sequencing libraries and cluster were established, primer was annealed and samples were sequenced on the PacBio RS II System as previously described ^34^. The obtained sequence raw data sets were analyzed to reveal the genome-wide base modification and the identification of m^6^A, m^4^C, and m^5^C of corresponding methylation (recognition) motifs. SMRT sequencing raw reads of low quality were filtered and the DNA fragment assembly was conducted. Reads were processed and mapped using BLAST mapper and the Pacific Biosciences SMRT Analysis pipeline using the standard mapping protocol. Reads from the strains were mapped to UTI89 genome reference sequence. The pulse width and inter pulse duration (IPD) ratio for each base were measured, which is arising when the DNA polymerase copies past the modification ^35^, and modification for each base was determined using an *in-silico* control. Sequence motif cluster analysis was done with a cutoff of −10log(*p-*value) score >30 (*P* value<0.001).

### RNA sequencing and DEGs analysis

Each of the strains used for RNA samples for three replicates was grown for 10h in LB medium, and then cell pellets were collected by centrifugation at 12,000 rpm (4□) to discard the supernatant. Total RNA was extracted using bacteria RNA kit (Omega Bio-tek, USA) according to manufacturer’s protocol. The RNA concentration was then determined using a Nanodrop 2000 machine and RNA quality was tested by visualization on agarose gels. Qualified total RNA was further purified and enriched. Following rRNA removal, fragmentation, synthesis of the second strand, adenylation of 3′ ends, adapter ligation, and amplification were performed before sequencing using an Illumina HiSeq 2500 (Illumina, San Diego, CA, USA). Reads were assessed and quantified before aligning to a reference genome (NCBI accession no. NC_007946.1) of *E. coli* UTI89. For the statistical analysis, using two criteria: |log_2_(Fold Change)|>2 and FDR < 0.01 as the cut-off values for screening significant differently expressed genes (DEGs). DEGs was performed using an EdgeR package in the R statistical environment, *P* values were adjusted for multiple testing with the Benjamini-Hochberg procedure as a false discovery rate (FDR) value ^36^.

### COG analysis

The predicted genes (corresponding to demethylated nucleotide sites) were derived from methylome and transcriptome in the *Δdam* compared to parent strain UTI89. COG categories and protein functional annotations for common DEGs were downloaded from the NCBI COG database ^37^. The amino acid sequences were aligned to the COG database as previously described ^38^. The COG IDs of all annotations were then classified based on their COG terms. Each pathway was matched to DEGs via in-house whole genome and transcripts.

### KEGG and GO pathway enrichment analysis

Pathways information from the KEGG (Kyoto encyclopedia of genes and genomes) database and GO (gene ontology) were used to investigate the response of biological processes to *dam* deletion. The top 10 significant enrichment of KEGG and GO were identified in the R statistical environment ^39^. We selected the KEGG pathways with a *P* value < 0.05 and the GO pathways with FDR < 0.05 as significantly enriched. A GO biological processes concept network plot (cnetplot) within the pathways was constructed. The enrichment information was visualized by the following: for KEGG pathways, it depicts enrichment scores using *P* values with different color and gene count as bar height, for GO pathways, the cnetplot displayed most significant terms and the linkages of genes involved.

### Comparative COG analysis of methylome and transcriptome of the *Δdam* mutant

COG database is a popular tool for comparative genome annotation. To better understand DEGs related protein function on a transcrioptomic level and allow tracing the effect of genomewide demethylation in *Δdam*, the transcriptome COG IDs were assigned in accordance with the cellular roles of the respective up- and down-regulated genes. An FDR value<0.05 is considered to be statistically significant. *P* values were adjusted for multiple testing with the Benjamini-Hochberg procedure as a false discovery rate (FDR) value ^36^. Data preparation and statistical analyis were performed using R software (R Foundation for Statistical Computing) and GraphPad Prism software (GraphPad).

## Results

### *dam* deletion mutation impaired persister formation in *E. coli* UTI89

To study the role of DNA methylation in *E. coli* persister formation, we constructed *dam* and *dcm* knockout mutants (*Δdam, Δdcm*) from the parent strain UTI89 and examined if the mutants had any defect in persister formation under various antibiotic exposure and stress conditions. We first analyzed the tolerance to diverse antibiotics (ampicillin, gentamicin, levofloxacin, ciprofloxacin, norfloxacin, cefalexin). Our results showed that the *Δdcm* mutation had no apparent effect on persister formation compared with UTI89 (data not shown). In contrast, the *Δdam* mutant had significant defect in persister formation with various antibiotic or stress exposures. Specifically, we found that the *Δdam* mutant was more easily killed by various antibiotics such that deletion of *dam* resulted in a ∼10^2^ fold decrease in persister formation. Although the *Δdam* mutant was initially killed to the same extent as the parent strain, during prolonged exposure to antibiotics, the *Δdam* mutant had diminished ability to form persisters in stationary phase (cultured for 10 h, ∼10^9^ CFU/mL) (Figure 1A-F), except for ampicillin, which is known to affect growing bacteria but is unable to kill nongrowing bacteria ^40^.

**Figure 1.**
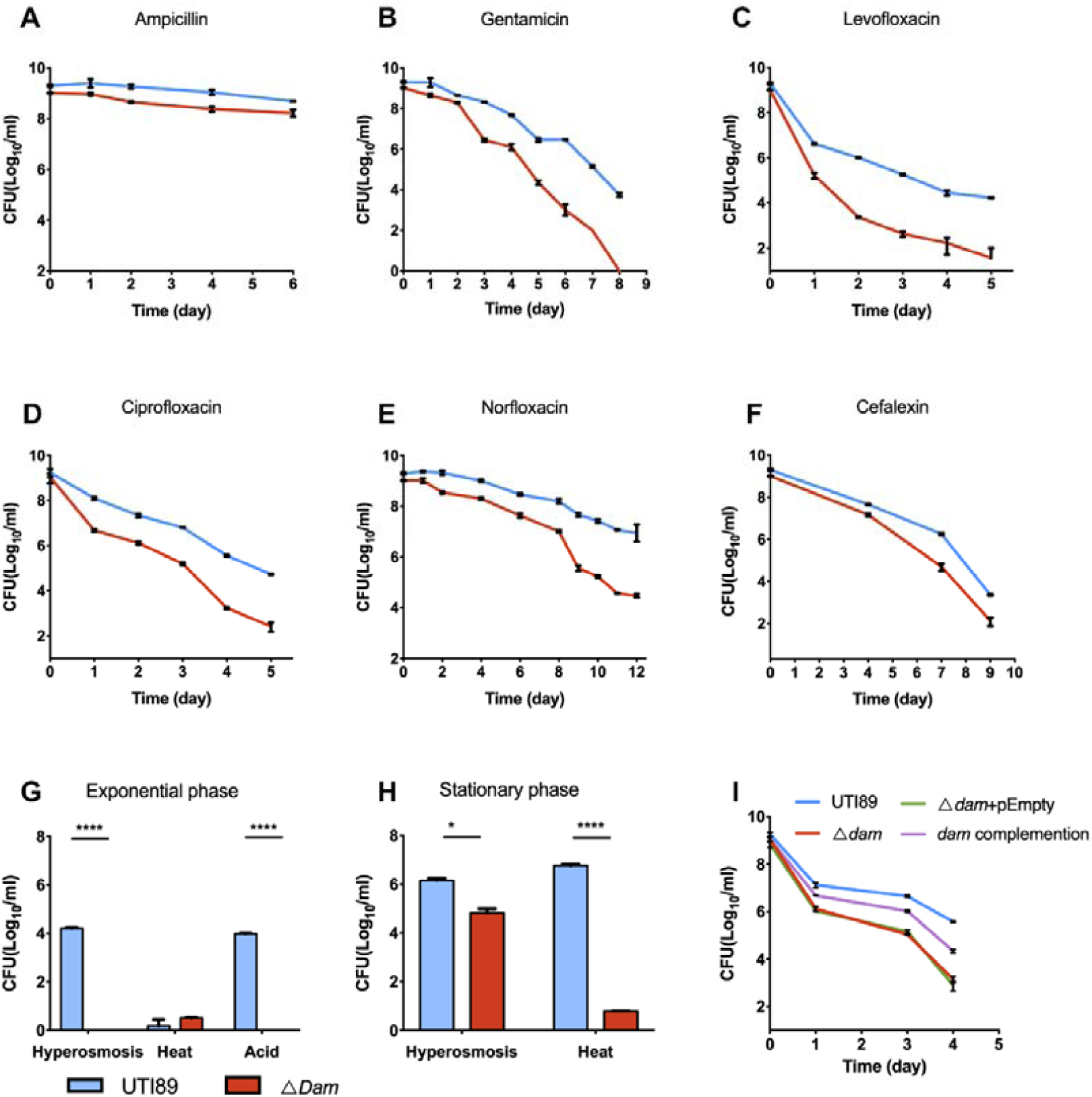
Persister formation of the *Δdam* mutant under various antibiotic treatment and multiple stresses. Persisters of stationary phase bacteria (∼10^9^ CFU/mL) were determined daily after exposure to (**A**) ampicillin (200 μg/ml), (**B**) gentamicin (40 μg/ml), (**C**) levofloxacin (10 μg/ml), (**D**) ciprofloxacin (10 μg/ml), (**E**) norfloxacin (8 μg/ml), (**F**) cefalexin (96 μg/ml). **(G).** Susceptibility of exponential phase bacteria (∼10^8^ CFU/mL) exposed to hyperosmosis (NaCl, 4 M) for 2d, heat (52□) for 1.5 h, and acid (pH 3.0) for 3d. (**H**) Susceptibility of stationary phase bacteria (∼10^9^ CFU/mL) exposed to hyperosmosis for 5d, heat (52□) for 1.5 h. (**I**) Persister formation of the *Δdam* mutant, *dam* complementation strain, *Δdam*+pEmpty vector and parent strain UTI89 under levofloxacin (7.5μg/ml) treatment.

Because persister formation was closely associated with cellular metabolic state and growth cycle, the age of bacterial culture could also affect the persister level ^41^. Thus, we evaluated effect of both log phase and stationary phase on the *Δdam* mutant in response to different stresses. The effects of stress on the *Δdam* mutant were determined in the exponential phase bacteria (cultured for about 3h, ∼10^8^ CFU/mL) with the following stresses: hyperosmosis (4M NaCl, 2d), heat (52□, 1.5h) and acid (pH 3.0, 3d). During the hyperosmosis and acid exposure, the *Δdam* mutant displayed a dramatic decrease in persister numbers compared with the parent strain UTI89, but no significant effect on sensitivity to the heat stress was seen (Figure 1G). When bacterial population was grown to stationary phase, non-growing bacteria were more tolerant to external stresses. Nevertheless, for stationary phase cultures, the *Δdam* mutant exhibited a similar defect in persister formation as shown by hypersensitivity to hyperosmosis (Figure 1H), as in exponential phase. In addition, complementation of the *Δdam* mutant restored the level of persisters to the wild-type level in antibiotic (levofloxacin) exposure assay, whereas the *Δdam* mutant transformed with empty vector remained susceptible as the *Δdam* mutant alone (Figure 1I).

### Genome-wide identification of Dam methylation

To identify the Dam methylation sites and targeted modification type, PacBio SMRT sequencing of whole genome DNA *de novo* was performed to determine the modification signature in the *Δdam* mutant compared with the parent strain UTI89 using the PacBio RS II System. The reads from the strains were mapped to the UTI89 genome reference sequence. The methylated motif (m^6^A, m^4^C and base modification) detected and the identification of corresponding methylation are shown in Table 1 (the *Δdam* mutant) and Table 2 (wild type UTI89). The motif string with methylated DNA base-modification sequences was detected, fraction referred to the proportion of relevant motif in the whole genome, and the mean score was used to evaluate the strength of modified base signal directly. Bacterial methylome can provide a wealth of information on the methylation marks present in bacterial genomes. The most obvious difference was that a total of 41359 genomic positions were found to be methylated in the parent strain UTI89, which were all demethylated in the *Δdam* mutant (demethylated motif strings in the *Δdam* mutant were marked in red in Table 2). Based on the base modification profiles, these methylated bases were found to be predominately m^6^A modifications of GATC (99.73%). DNA adenine methylase methylates the adenine in GATC site, and the methylome analysis indicated that almost all the GATC motifs in the genomic DNA of the *Δdam* mutant were demethylated because of the *dam* deletion.

**Table 1.**
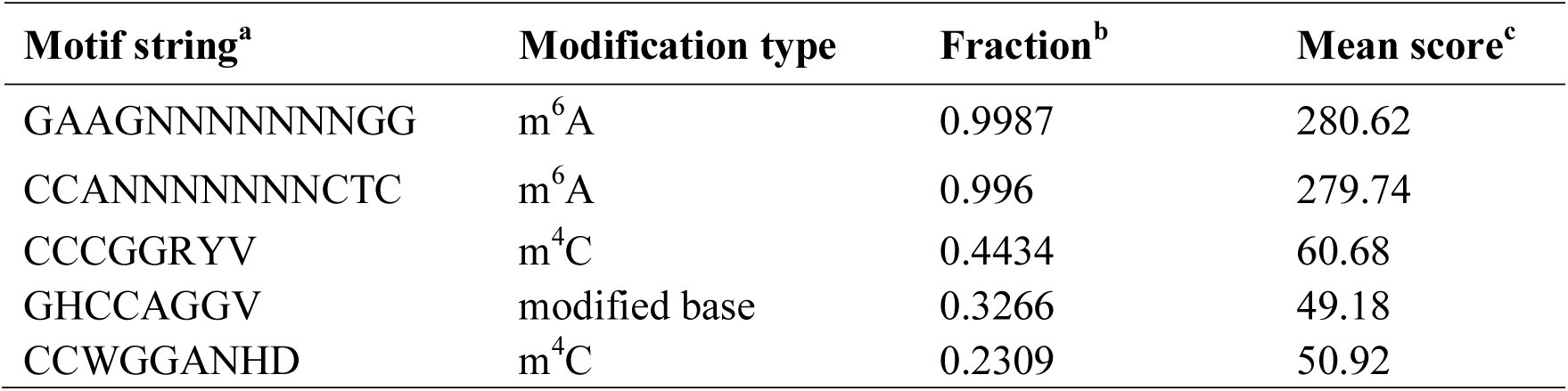
DNA base modifications in the *Δdam* mutant.

**Table 2.** DNA base modifications in wild type UTI89.

| Motif string<sup>a</sup> | Modification type | Fraction<sup>b</sup> | Mean score<sup>c</sup> |
| --- | --- | --- | --- |
| GAAGNNNNNNNTGG | m<sup>6</sup>A | 0.9987 | 237.40 |
| CCANNNNNNNCTTC | m<sup>6</sup>A | 0.996 | 239.97 |
| **GATC** | **m<sup>6</sup>A** | **0.9973** | **253.76** |
| CCCGGRY | m<sup>4</sup>C | 0.4965 | 65.61 |
| **CCAGGAH** | **m<sup>4</sup>C** | **0.4229** | **52.58** |
| **TCCGGRYAD** | **modified base** | **0.3577** | **49.14** |
| GHCCAGGR | modified base | 0.3405 | 49.78 |
| CCWGGVNYR | modified base | 0.1721 | 46.88 |

<sup>a</sup>N: A/T/C/G, R: A/G, Y: C/T, V: G/A/C, H: A/T/C, D: G/A/T, W: A/T
<sup>b</sup>Fraction: proportion of relevant motif in the whole genome.
<sup>c</sup>Mean score: to evaluate the strength of modified base signal directly.

The genomewide distribution of methylated bases in UTI89 and the *Δdam* mutant was determined using PacBio SMRT sequencing technology. Clustering of methylated nucleotides based on sequence context identified 8 recognition motifs in the parent strain UTI89, which were m^6^A, m^4^C and modified base methylation (Figure 2A). In the *Δdam* mutant, 5 recognition motifs were determined, however, 3 motifs matched to UTI89 were not detected, of which, the main methylation of recognition motif is GATC (m^6^A) (Figure 2B, red arrow in 2A) compared with UTI89. The results showed that adenine of GATC in the *Δdam* mutant was demethylated. Different methylation types would affect the gene expression and relevant protein function. To better understand the global responses to demethylation, we performed a comparative analysis between the differentially methylated sites related genome-informatics in UTI89 and the *Δdam* mutant (Figure 2C, D). Common methylation motif related genes were classified through protein functions determined from the COG database. Classification of predicted proteins is highlighted with different colors in the whole genome, of which, the circles are GC level, GC skew, RNA, positive-strand COG and negative-strand COG in UTI89 (Figure 2C) and the *Δdam* mutant (Figure 2D) from inside to outside. Differentially methylated sites in *Δdam* are shown in Figure 2E. COG analysis assigned 4296 genes to 22 functional categories (Table 3), with the top 10 including the following categories: general function prediction only (R, 10.59%), carbohydrate transport and metabolism (G, 9.71%), function unknown (S, 8.99%), amino acid transport and metabolism (E, 8.38%), transcription (K, 7.36%), energy production and conversion (C, 6.98%), cell wall/membrane/envelope biogenesis (M, 6.03%), inorganic ion transport and metabolism (P, 5.66%), and ribosomal structure and biogenesis (J, 4.38%). COG analysis of the relevant gene clusters suggested the overwhelming effects caused by defects in methylation of GATC motifs on the genome-wide transcription involved in diverse cellular functions such as metabolism, transcription, translation, DNA repair, energy production, signal transduction, cell motility, and transport pathways (Table 3).

**Table 3.** COG class and distribution of predicted protein function of demethylated genes in *Δdam* versus UTI89.

| Function annotation: COG class | Count | Percentage |
| --- | --- | --- |
| R: General function prediction only | 455 | 10.59% |
| G: Carbohydrate transport and metabolism | 417 | 9.71% |
| S: Function unknown | 386 | 8.99% |
| E: Amino acid transport and metabolism | 360 | 8.38% |
| K: Transcription | 316 | 7.36% |
| C: Energy production and conversion | 300 | 6.98% |
| M: Cell wall/membrane/envelope biogenesis | 259 | 6.03% |
| P: Inorganic ion transport and metabolism | 243 | 5.66% |
| L: Replication, recombination and repair | 209 | 4.86% |
| J: Translation, ribosomal structure and biogenesis | 188 | 4.38% |
| T: Signal transduction mechanisms | 188 | 4.38% |
| H: Coenzyme transport and metabolism | 164 | 3.82% |
| O: Posttranslational modification, protein turnover, chaperones | 155 | 3.61% |
| U: Intracellular trafficking, secretion, and vesicular transport | 151 | 3.51% |
| N: Cell motility | 124 | 2.89% |
| I: Lipid transport and metabolism | 104 | 2.42% |
| F: Nucleotide transport and metabolism | 99 | 2.30% |
| Q: Secondary metabolites biosynthesis, transport and catabolism | 82 | 1.91% |
| V: Defense mechanisms | 57 | 1.33% |
| D: Cell cycle control, cell division, chromosome partitioning | 35 | 0.81% |
| A: RNA processing and modification | 2 | 0.05% |
| W: Extracellular structures | 2 | 0.05% |

**Figure 2.**
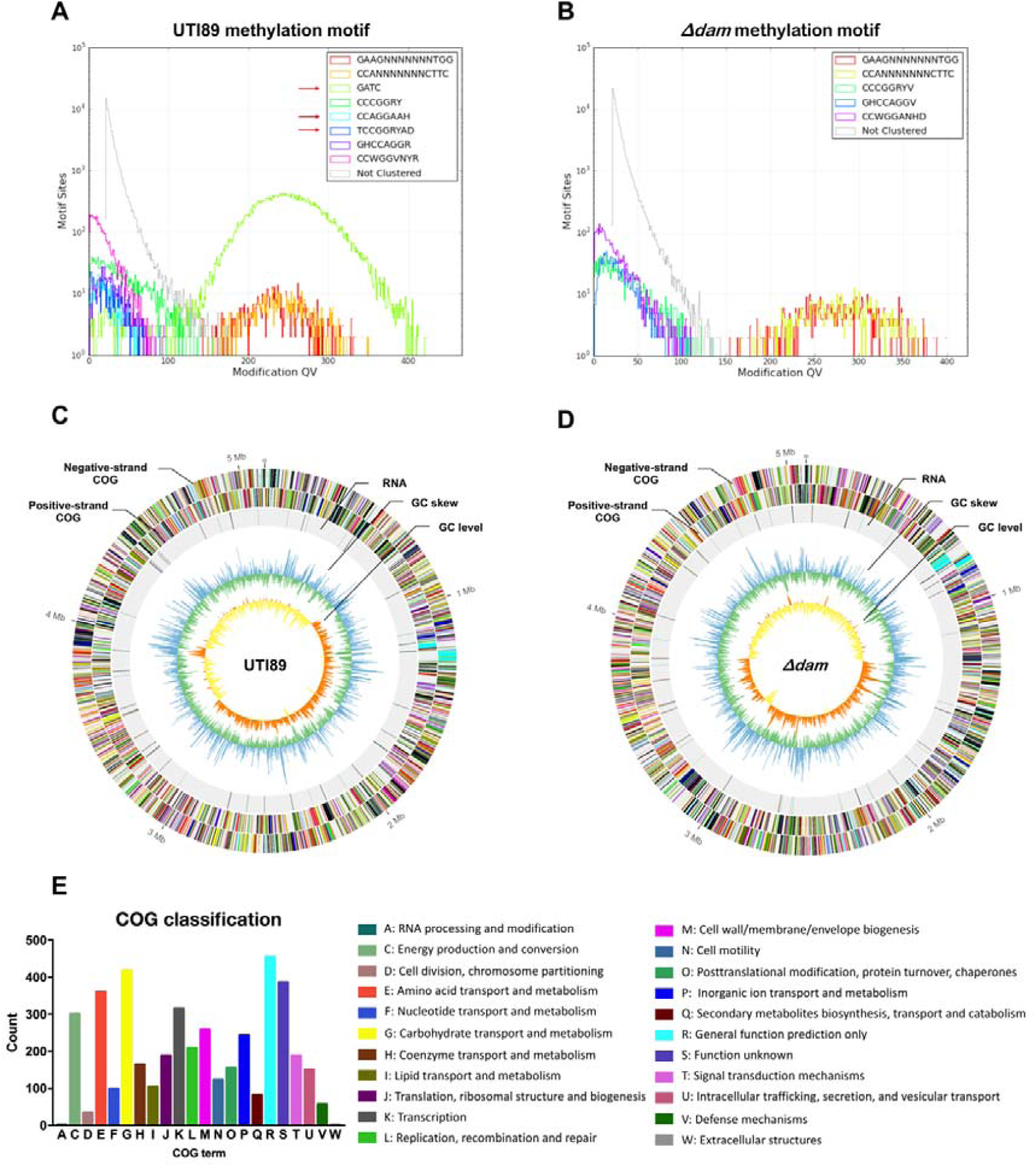
Comparison of UTI89 and *Δdam* methylome. (A) The methylation patterns and corresponding motifs detected in UTI89. (B) The methylation patterns and corresponding motifs detected in *Δdam*. (C) Circle map displaying the distribution of methylated bases in UTI89 chromosome, including RNA, GC characters and COG classification of detected methylation-site associated genes. (D) Circle map displaying the distribution of methylated bases in *Δdam* chromosome, including RNA, GC characters and COG classification of detected methylation-site associated genes. Tick marks display the genomic positions in megabases. (E) COG classification and distribution of predicted protein function of demethylated nucleotide site related genes in *Δdam*. COG terms are highlighted with different colors.

### Comparative transcriptome analysis of the *Δdam* mutant

In addition to global characterization of methylation patterns in genomic DNA and function annotations, we sought to focus on the effect of demethylated GATC sites on the transcription of the *Δdam* mutant, to shed light on its effect on altering vital pathways of persister formation. To this end, we generated RNA sequencing (RNA-seq) data of UTI89 and *Δdam*. With both |log_2_Fold Change| >2 and false discovery rate (FDR) < 0.01 as cut-off criteria, the total number of DEGs is listed in Supplementary Table 2. Comparison of *Δdam* to the parent strain UTI89 identified 1345 differentially expressed genes (DEGs). Of these DEGs, 153 genes were significantly upregulated and 1192 genes were significantly downregulated (Figure 3B). Heatmap cluster analysis of these selected genes revealed that the expression profiles of the control and the *Δdam* mutant were distinct, and the set of genes significantly down-regulated in *Δdam* but not in UTI89 (Figure 3A), is of especial interest, as these genes may be involved in persister formation, such as *fnr, usp* operons, *fim* clusters, *tra* operons, *repA, ccdAB* (type II toxin-antitoxin system toxin), *tss* clusters (type VI secretion system), *arcAC, gsp* clusters (type II secretion system), GntR family transcriptional regulator (Supplementary Table 2), some of which are known to be involved in persister formation. Functional KEGG enrichment analysis of the DEGs was identified, the top 10 most significantly enriched biological processes (up and down DEGs) are listed in Figure 3C and Supplementary Table 3. The largest number of upregulated DEGs were enriched in flagellar assembly (eco02040) in the *Δdam* mutant, while the down-regulated DEGs were significantly enriched in ABC transporter (eci02010) pathway. In addition, compared with UTI89, galactose metabolism and propanoate metabolism were overexpressed, whereas transmembrane transport sugar phosphotransferase systems (PTS) and secretion-related pathways were down-regulated in the *Δdam* mutant.

**Figure 3.**
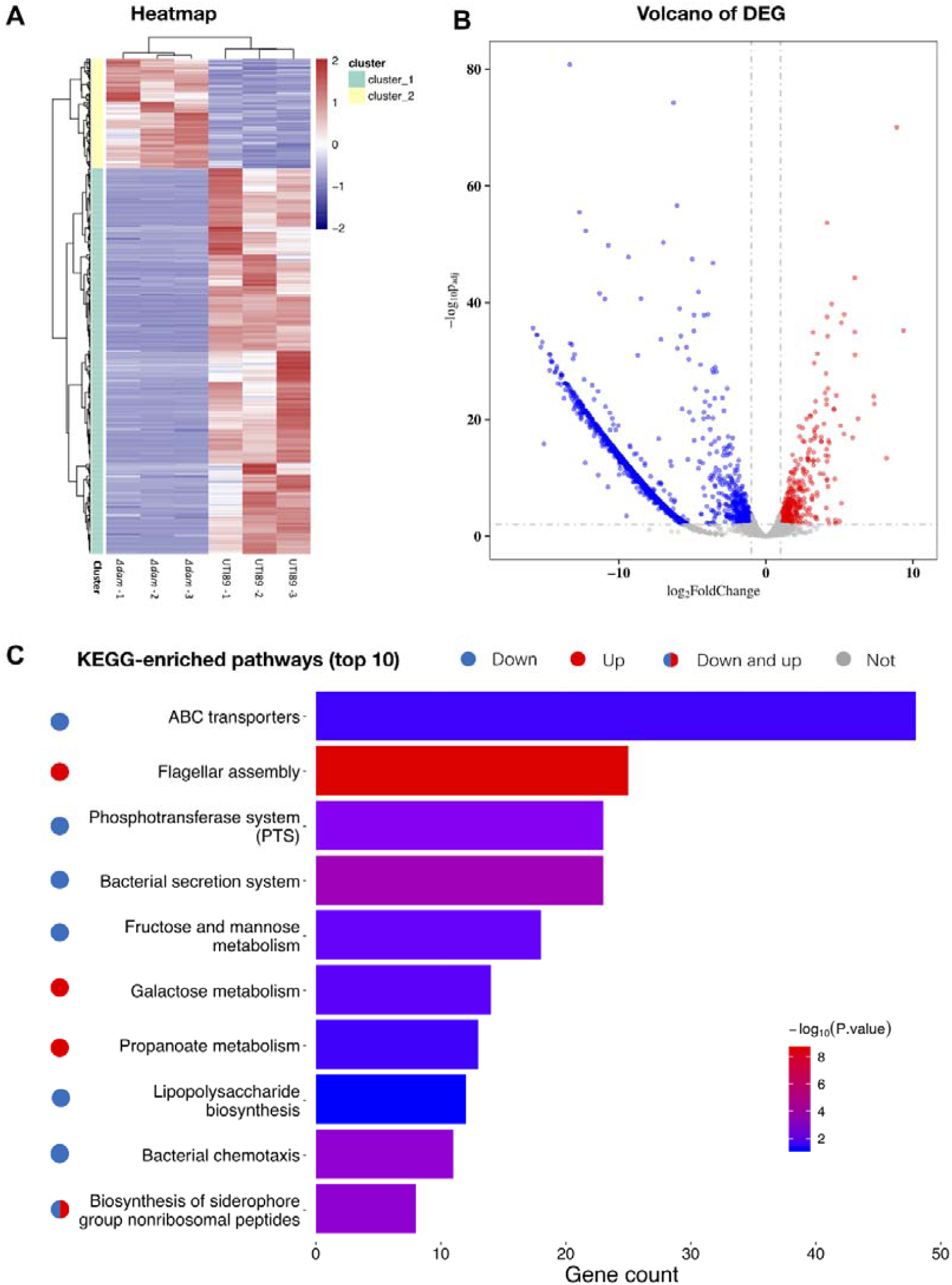
RNA-seq analysis and KEGG pathway of DEGs in the *Δdam* mutant. (A) Heatmap showing clear differences between *Δdam* and UTI89 in terms of relative levels of expression (upregulation, red) or (downregulation, blue). Row values were calculated using gene expression levels (in fragments per kilobase per million; FPKM). (B) Volcano plots of DEGs between the *Δdam* mutant and UTI89.. In plots, circle points indicate each gene of *Δdam*. Two criteria of |log_2_(fold change)| >2 and false discovery rate (FDR) < 0.01 are highlighted in dotted lines, respectively. (C) Top 10 enriched KEGG pathways of DEGs. Enrichment analysis was performed with a significant criterion, *P* value < 0.05. The regions were ranked by *P* value. Log_10_(*P* value) ranged from blue (relatively low) to red (relatively high) in the *Δdam* mutant versus UTI89. For each KEGG pathway, the bar shows the number of enriched genes. The red circle points indicate significantly upregulated, the blue circle points indicate significantly downregulated DEGs, the red + blue circle points indicate significantly up and down regulated DEGs and the gray circle points indicate no statistical significance.

### Functional analysis by biological network to identify key pathways in the *Δdam* mutant

To compare functional differences between UTI89 and *Δdam*, we used an FDR of 0.05 to perform a GO term enrichment analysis and performed cnetplot (concept network plot) analysis based on DEGs (Figure 4). It shows cluster distribution and relationship of proteins in each pathways. We identified enriched pathways in upregulated and downregulated genes from the *Δdam* mutant compared to the parent strain UTI89 (Figure 4). The upregulated pathways include translation (GO:0006412) and flagellum-dependent cell motility (GO:0071973), and downregulated processes include cell adhesion (GO:0007155) (fimbriae relevant protein), DNA integration (GO:0015074) (transposase, integrase and recombinase), DNA recombination and repair (GO:0006310) (integrase, excisionase, recombinase and endodeoxyribonuclease) and cytolysis (GO:0019835) (lysis protein) (Supplementary Table 4).

**Figure 4.**
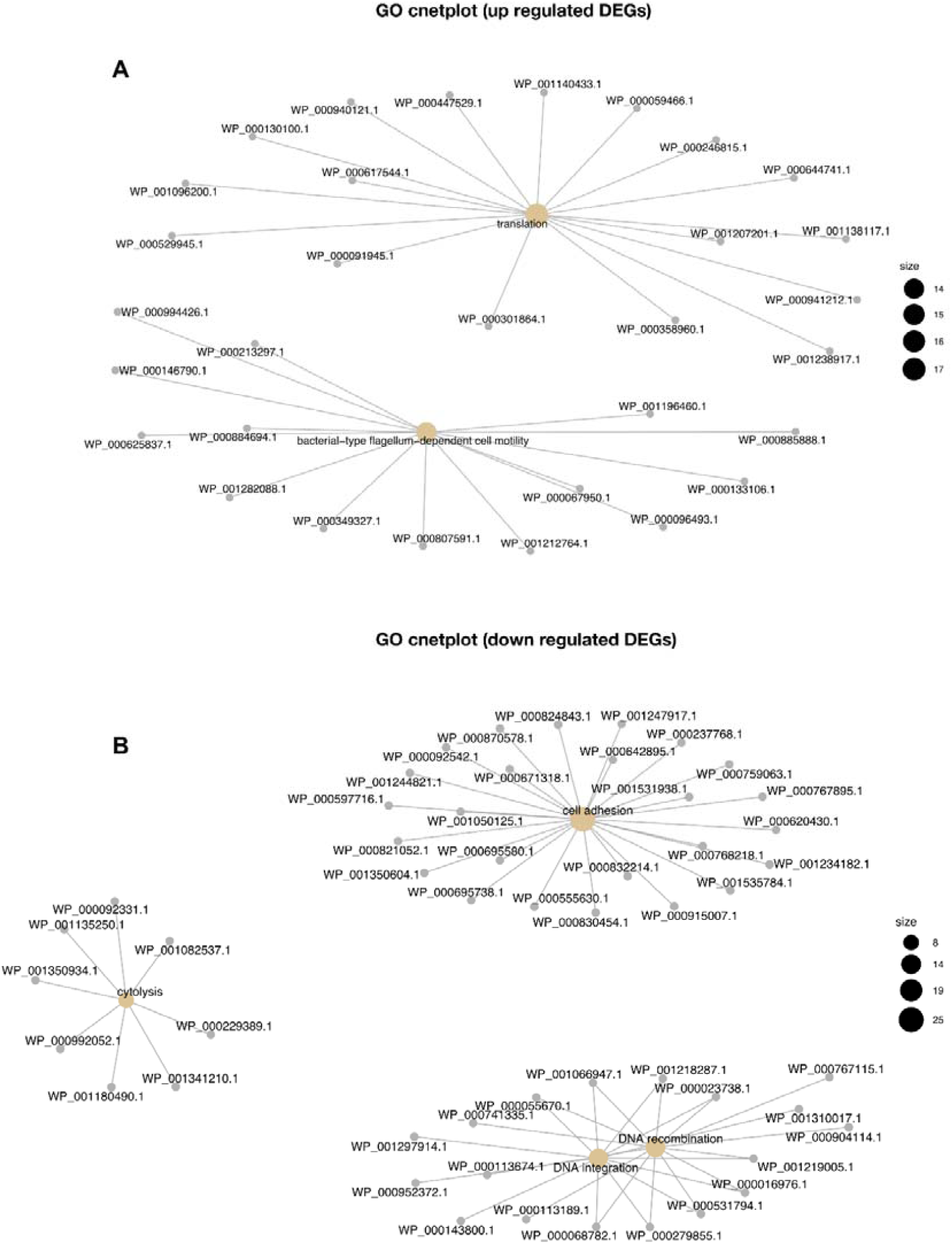
Gene ontology term enrichment cnetplot analysis of DEGs in *Δdam*. (A) Biological processes enriched from up-regulated DEGs. (B) Biological processes enriched from down-regulated DEGs. The brown nodes represent key pathways involved based on regulated DEGs and predicted protein functions. The size of the nodes reflects the number of enriched DEGs. Enrichment analysis was performed with a significant FDR < 0.05.

### Comparative analysis of transcriptome and methylome COG class in *Δdam*

To further verify the alteration of gene expression that is related to demethylation in *Δdam*, we performed COG functional category analysis of transcriptome and compared with COG terms of demethylated annotation (Figure 5A). The trancriptome COG demonstrated a correlation with predicted methylome COG, and the results showed that up-regulated DEGs accounted for a relatively small fraction, with statistically significant (FDR<0.05) pathways mapped in J (translation, ribosomal structure and biogenesis) and N (cell motility) being identified (Figure 5B). Most of genes were down-regulated and annotated to multiple functional catagories that were extensively affected by the DNA demethylation, with statistically significant (FDR<0.05) pathways mapped in L (DNA replication, recombination and repair) and Q (secondary metabolite biosynthesis, transport and catabolism) being identified (Figure 5C). These alterations could be associated with diminished persister formation in *Δdam*.

**Figure 5.**
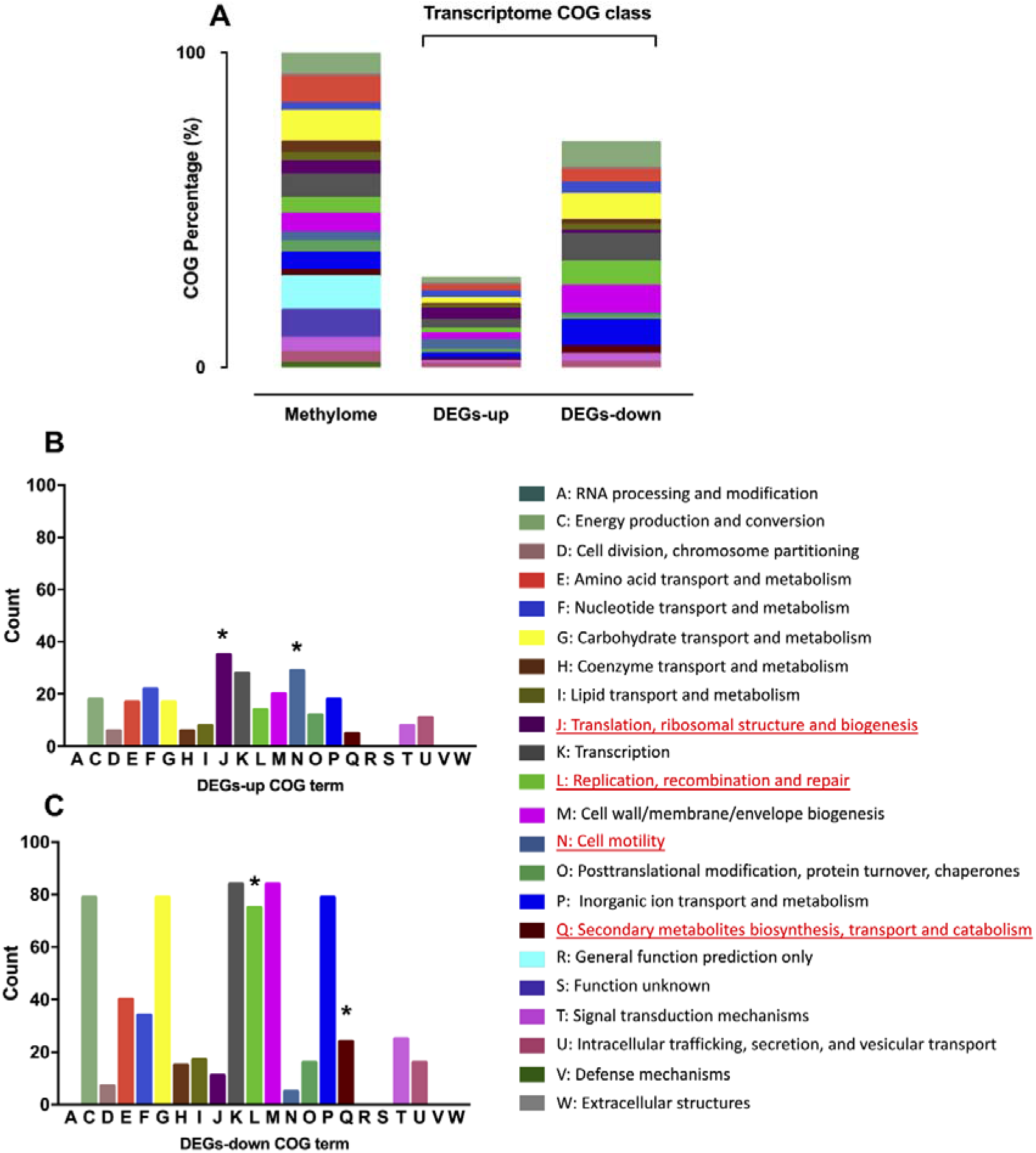
The comparative analysis of transcriptome and methylome COG in *Δdam* compared with UTI89. **(A)** The transcriptome COG IDs are mapped to methylome COG classification. COG terms of transcritps are assigned by the cellular function of respective up-regulated **(B)** and down-regulated **(C)** genes (*FDR<0.05). The underlined and red marked terms are significant functional categories.

## Discussion

Although DNA adenine methylation in bacterial genome has been shown to affect various biological processes such as transcriptional control, mismatch repair, DNA replication, and host-pathogen interactions in bacteria ^19, 21^, the link between DNA methylation and bacterial persister formation has not been investigated previously. In this study, we found that the *Δdam* mutant but not the *Δdcm* mutant of *E. coli* UTI89 had significant defect in persistence to multiple antibiotics and stresses compared with the *dam* (+) parent strain. In addition, complementation of the *Δdam* mutant with the wild type gene restored the persistence phenotype close to the wild type level, which confirms the role of DNA adenine methylation in persister formation. Although a previous study examined the transcription of a small number of targeted genes (*papIB, papEF, qnrA, arcA, gyrB, mdh, recA, and rpoS*) in a *dam* mutant in different UPEC strains *E. coli* C119 and CFT073 ^42^, here we performed a more comprehensive whole genome and transcriptome level analysis by SMRT and RNA-seq and obtained crucial methylome information linking the gene expression differences in the *Δdam* mutant strain which provides important insights and potential explanation about its defect in persistence. It is of interest to note that the methylated state is usually correlated with transcriptional repression ^19^, and loss of methylation due to *dam* mutation could lead to a more generalized transcriptional activation of genes in the *Δdam* mutant, which could partly explain the reduced persister formation of the *Δdam* mutant.

Our genome-wide methylome analysis of the *Δdam* and UTI89 showed that the most obvious difference was that there existed 41359 m^6^A modifications of GATC in UTI89 which were all demethylated in the *Δdam* mutant (Table 1, 2). The disparity illustrated that the deletion of *dam* in UTI89 caused an overwhelming effect on the whole genome (4296 genes of relevant predicted marks), and the distribution of COG classification analysis identified many genes involved in key pathways including metabolism, transmembrane process, transcription, cell motility being affected (Table 3). Lacey et al. ^43^ conducted Dam methylation analysis of different growth phases (log phase, stationary phase, death phase) of *E. coli* and indicated a role of Dam in transcription and cell survival. It is interesting to note that genes *fadR, parE* involved in transmembrane transport/nucleoside binding/sulfur metabolism overlapped in our study. In addition, many of methylated sites identified in the above study of the wild type strain of stationary phase are located within genes that encode proteins involved in transport or transcriptional regulation, which were downregulated in the *Δdam* mutant in our study, a finding that is consistent with their observations.

Our transcriptome analysis identified that crucial transcriptional regulators were massively downregulated in *Δdam* (Supplementary Table 2), these regulators are known to be involved in persister formation with the actication of stress response, including LexA^44^ NtrB ^45^, Fis ^46^, Fnr, Lrp ^47^, SoxS ^48^, ArcA ^47 49^, which may be responsible for the defect in persister formation in the *Δdam* mutant or survival in various stresses as shown in this study. It also revealed that demethylation in the *Δdam* mutant impaired persister formation by affecting extensive gene expression, such as upregulating genes involved in flagellar synthesis and assembly and galactose metabolism, propanoate metabolism and downregulating genes involved in different types of fimbrial biosynthesis, pilus synthesis, metabolism (carbon and nitrogen source), sugar phosphotransferase systems (PTS) and secretion-related pathways.

The transcriptome data analysis indicated that Dam tightly controls flagella transcription in *E. coli*, as shown by upregulation the expression of flagella biosynthesis including flagella biosynthesis (*flg* operon) and assembly (*fli* operon) process in the *Δdam* mutant. The results were consistent with previous observations that flagella synthesis genes (*flgE, flgJ, fliG, flhB*) were related to persister formation ^41, 50^. It is likely that upregulation of flagella genes in the *Δdam* mutant is a reflection of suppression of flagella genes by Dam methylation in normal wild type bacteria which is important for persister formation. The overexpression of the flagella genes in the *Δdam* mutant is reminiscent of the PhoU mutant ^41^, where PhoU serves as a metabolic switch in suppressing cellular metabolism in the wild type strain but its loss of function in the mutant leads to hyperactive metabolism and inability to form persisters thus showing higher susceptibility to antibiotics and stresses. Morever, we observed the most enriched KEGG and GO pathways in persister formation of *Δdam* mutant include cell adhesion, motility and translation. Previous studies have shown virulence attenuation in *Salmonella* Dam mutants with defective DNA methylation ^21, 51^, since UPEC surface appendages such as fimbria and pili are known virulence factors involved in attachment and adhesion to bladder epithelial cells, the downregulation of genes for these structures as seen in the *Δdam* mutant in this study is consistent with the observation that DNA methylation is important for virulence of UPEC bacteria as mutations in DNA methylation caused virulence attenuation and defect in biofilm formation ^52^. Extensive pili operons involved in adhesion and virulence in UPEC including *sfa, fimHX, auf, pap* and *tra* related fimbriae were down-regulated in the *Δdam* mutant and indicate their regulation by DNA methylation. Previous studies described that methylation of the *pef* GATC II site by Dam were required for plasmid-encoded fimbriae (*pef*) transcription ^53^, pyelonephritis-associated pili (*pap*) promoter was governed by DNA methylation state in GATC sites, expression of *pap* pili was controlled by phase variation (cell cycle), resulting in the heterogeneous bacterial population with pili (“phase ON” state) or without pili (“phase” OFF state) ^54^. It will be of interest to determine if the *Δdam* mutant is more attenuated for virulence and persistence in a UTI animal model in future studies. Dam acts as a switch for global gene regulation, and previous studies showed the connection between DNA methylation and stationary phase transcription ^43, 55^, and our finding that DNA methylation is more specifically involved in persister formation is an extension of this observation. Since Dam in *E. coli* is necessary for cell cycle timing through methylation of *oriC* ^56^, future studies can be performed to assess the role of Dam methylation in cell cycle regulation in possible transition of growing cells to persister cells.

Furthermore, our comparative analysis of omics COG data identified potentially the most important pathways responsible for persister formation due to the altered methylation nucleotide sites (Figure 5A). We found that J (translation, ribosomal structure and biogenesis), N (cell motility) (Figure 5B) were significantly up-regulated and L (DNA replication, recombination and repair) and Q (secondary metabolites biosynthesis, transport and catabolism) (Figure 5C) were significantly down-regulated. It is known that DNA replication and repair are essential to DNA damage response, and persister cells require DNA repair machinery (RecA, RecB) to survive the antibiotic stress ^57 50^, and it is of interest to note that *recA* involved in DNA repair pathway was downregulated in the *Δdam* mutant in our study. In addition, the sencondary metabolites were reported to be important for bacterial sensing and damage response to the adverse environment, for secondary metabolites achieving ther functions, ATP-binding cassette (ABC) transmembrane transporters are crucial ^58^. The upregulation of genes in translation, motility and the downregulation of genes in DNA repair, transport due to *dam* deletion provides possible explanation of significantly diminished persister formation. Taken together, the COG classification analyses are consistent with the KEGG and GO functional annotation results and suggest that Dam mediates persister formation through affecting multiple pathways including translation, motility, signaling and transport, DNA repair in the cell.

In summary, we found that the *Δdam* mutant had a significant defect in persister formation under antibiotic and stress conditions. Our findings suggest that DNA adenine methylation plays an important role in persister formation by maintaining genome stability (DNA replication, recombination and repair) and systematically affecting diverse cellular functions including bacterial adhesion (fimbriae and pilus), flagella biosynthesis, galactitol transport/utilization, diverse transport systems, metabolism (carbon, purine, and amino acid) and transcriptional repression. Further studies are necessary to determine DNA methylation specific switch factor on the crucial transition process toward persister formation. In addition, it would be of interest to determine if the *Δdam* mutant has attenuated virulence and defect in creating persistent infection in animal models in the future. Because of the importance of Dam in persister formation, we believe Dam may serve as a novel therapeutic target for future drug development against bacterial persisters.

## Acknowledgement

This work was supported by National Science Foundation of China (81572046) (81772231).

## Author contributions

Y.Z., W.H.Z conceived and designed the experiments, W.H.Z supervised the work, S.L., Y.Y.X. performed the experiments, S.L., Y.Y.X., Y.Z. analyzed the results. Y.Y.X. wrote the manuscript.

## Competing interests

The authors declare no competing interests.

## References

1. Foxman B. Epidemiology of urinary tract infections: incidence, morbidity, and economic costs. Am J Med 113 Suppl 1A, 5s–13s (2002).

2. Flores-Mireles AL, Walker JN, Caparon M, Hultgren SJ. Urinary tract infections: epidemiology, mechanisms of infection and treatment options. Nat Rev Microbiol 13, 269–284 (2015).

3. Kostakioti M, Hultgren SJ, Hadjifrangiskou M. Molecular blueprint of uropathogenic Escherichia coli virulence provides clues toward the development of anti-virulence therapeutics. Virulence 3, 592–594 (2012).

4. Juan J. Martinez MAM, Joel D. Schilling, Jerome S. Pinkner, Scott J. Hultgren. Type 1 pilus-mediated bacterial invasion of bladder epithelial cells. EMBO J 19, 2803–2812 (2000).

5. Ejrnaes K, et al. Pulsed-field gel electrophoresis typing of Escherichia coli strains from samples collected before and after pivmecillinam or placebo treatment of uncomplicated community-acquired urinary tract infection in women. J Clin Microbiol 44, 1776–1781 (2006).

6. Jantunen ME, Saxen H, Salo E, Siitonen A. Recurrent urinary tract infections in infancy: relapses or reinfections? J Infect Dis 185, 375–379 (2002).

7. Koljalg S, Truusalu K, Vainumae I, Stsepetova J, Sepp E, Mikelsaar M. Persistence of Escherichia coli clones and phenotypic and genotypic antibiotic resistance in recurrent urinary tract infections in childhood. J Clin Microbiol 47, 99–105 (2009).

8. Luo Y, et al. Similarity and divergence of phylogenies, antimicrobial susceptibilities, and virulence factor profiles of Escherichia coli isolates causing recurrent urinary tract infections that persist or result from reinfection. J Clin Microbiol 50, 4002–4007 (2012).

9. Bigger J. Treatment of staphylococcal infections with penicillin by intermittent sterilisation. The Lancet 244, 497–500 (1944).

10. Barber AE, Norton JP, Spivak AM, Mulvey MA. Urinary tract infections: current and emerging management strategies. Clin Infect Dis 57, 719–724 (2013).

11. Frimodt-Moller J, Lobner-Olesen A. Efflux-Pump Upregulation: From Tolerance to High-level Antibiotic Resistance? Trends Microbiol 27, 291–293 (2019).

12. Fluit AC, Schmitz FJ. Bacterial resistance in urinary tract infections: how to stem the tide. Expert Opin Pharmacother 2, 813–818 (2001).

13. Nira Rabin YZ, Clement Opoku-Temeng, Yixuan Du, Eric Bonsu, Herman O Sintim. Biofilm formation mechanisms and targets for developing antibiofilm agents. Future Med Chem 7, 493–512 (2015).

14. Aseev LV, Koledinskaya LS, Boni IV. Regulation of Ribosomal Protein Operons rplM-rpsI, rpmB-rpmG, and rplU-rpmA at the Transcriptional and Translational Levels. J Bacteriol 198, 2494–2502 (2016).

15. Kaspy I, Rotem E, Weiss N, Ronin I, Balaban NQ, Glaser G. HipA-mediated antibiotic persistence via phosphorylation of the glutamyl-tRNA-synthetase. Nat Commun 4, 3001 (2013).

16. Amato SM, et al. The role of metabolism in bacterial persistence. Front Microbiol 5, 70 (2014).

17. Zhang Y. Persisters, persistent infections and the Yin-Yang model. Emerging microbes & infections 3, e3 (2014).

18. Van den Bergh B, Fauvart M, Michiels J. Formation, physiology, ecology, evolution and clinical importance of bacterial persisters. FEMS Microbiol Rev 41, 219–251 (2017).

19. Casadesus J, Low D. Epigenetic gene regulation in the bacterial world. Microbiol Mol Biol Rev 70, 830–856 (2006).

20. Marinus MG MN. Isolation of deoxyribonucleic acid methylase mutants of Escherichia coli K-12. J Bacteriol 114, 1143–1150 (1973).

21. Erova TE, Kosykh VG, Sha J, Chopra AK. DNA adenine methyltransferase (Dam) controls the expression of the cytotoxic enterotoxin (act) gene of Aeromonas hydrophila via tRNA modifying enzyme-glucose-inhibited division protein (GidA). Gene 498, 280–287 (2012).

22. Wion D, Casadesús J. N6-methyl-adenine: an epigenetic signal for DNA-protein interactions. Nat Rev Microbiol 4, 183–192 (2006).

23. Marinus MG, Casadesus J. Roles of DNA adenine methylation in host-pathogen interactions: mismatch repair, transcriptional regulation, and more. FEMS Microbiol Rev 33, 488–503 (2009).

24. Niller HH, Masa R, Venkei A, Meszaros S, Minarovits J. Pathogenic mechanisms of intracellular bacteria. Curr Opin Infect Dis 30, 309–315 (2017).

25. Ugur S, Akcelik N, Yuksel FN, Taskale Karatug N, Akcelik M. Effects of dam and seqA genes on biofilm and pellicle formation in Salmonella. Pathog Glob Health 112, 368–377 (2018).

26. Aya Castaneda Mdel R, Sarnacki SH, Noto Llana M, Lopez Guerra AG, Giacomodonato MN, Cerquetti MC. Dam methylation is required for efficient biofilm production in Salmonella enterica serovar Enteritidis. Int J Food Microbiol 193, 15–22 (2015).

27. Costa KC, Glasser NR, Conway SJ, Newman DK. Pyocyanin degradation by a tautomerizing demethylase inhibits Pseudomonas aeruginosa biofilms. Science 355, 170–173 (2017).

28. Atack JM, et al. A biphasic epigenetic switch controls immunoevasion, virulence and niche adaptation in non-typeable Haemophilus influenzae. Nat Commun 6, 7828 (2015).

29. Stewart PS. Mechanisms of antibiotic resistance in bacterial biofilms. Int J Med Microbiol 292, 107–113 (2002).

30. Datsenko KA, Wanner BL. One-step inactivation of chromosomal genes in Escherichia coli K-12 using PCR products. Proc Natl Acad Sci U S A 97, 6640–6645 (2000).

31. Liu S, Wu N, Zhang S, Yuan Y, Zhang W, Zhang Y. Variable Persister Gene Interactions with (p)ppGpp for Persister Formation in Escherichia coli. Front Microbiol 8, 1795 (2017).

32. Ma C, Sim S, Shi W, Du L, Xing D, Zhang Y. Energy production genes sucB and ubiF are involved in persister survival and tolerance to multiple antibiotics and stresses in Escherichia coli. FEMS Microbiol Lett 303, 33–40 (2010).

33. Zhang S, Liu S, Wu N, Yuan Y, Zhang W, Zhang Y. Small Non-coding RNA RyhB Mediates Persistence to Multiple Antibiotics and Stresses in Uropathogenic Escherichia coli by Reducing Cellular Metabolism. Front Microbiol 9, 136 (2018).

34. Fang G, et al. Genome-wide mapping of methylated adenine residues in pathogenic Escherichia coli using single-molecule real-time sequencing. Nat Biotechnol 30, 1232–1239 (2012).

35. Flusberg BA, et al. Direct detection of DNA methylation during single-molecule, real-time sequencing. Nat Methods 7, 461–465 (2010).

36. Benjamini Y, Hochberg Y. Controlling The False Discovery Rate - A Practical And Powerful Approach To Multiple Testing. J Royal Statist Soc, Series B 57, 289–300 (1995).

37. Galperin MY, Makarova KS, Wolf YI, Koonin EV. Expanded microbial genome coverage and improved protein family annotation in the COG database. Nucleic Acids Res 43, D261–269 (2015).

38. Buchfink B, Xie C, Huson DH. Fast and sensitive protein alignment using DIAMOND. Nat Methods 12, 59–60 (2015).

39. Qureshi R, Sacan A. Weighted set enrichment of gene expression data. BMC Syst Biol 7 Suppl 4, S10 (2013).

40. Tuomanen E, Cozens R, Tosch W, Zak O, Tomasz A. The rate of killing of Escherichia coli by beta-lactam antibiotics is strictly proportional to the rate of bacterial growth. J Gen Microbiol 132, 1297–1304 (1986).

41. Li Y, Zhang Y. PhoU is a persistence switch involved in persister formation and tolerance to multiple antibiotics and stresses in Escherichia coli. Antimicrob Agents Chemother 51, 2092–2099 (2007).

42. Stephenson SA-M, Brown PD. Epigenetic Influence of Dam Methylation on Gene Expression and Attachment in Uropathogenic Escherichia coli. Front Public Health 4, 131 (2016).

43. Westphal LL, Sauvey P, Champion MM, Ehrenreich IM, Finkel SE. Genomewide Dam Methylation in Escherichia coli during Long-Term Stationary Phase. mSystems 1, e00130–00116 (2016).

44. Leung V, Ajdic D, Koyanagi S, Levesque CM. The formation of Streptococcus mutans persisters induced by the quorum-sensing peptide pheromone is affected by the LexA regulator. J Bacteriol 197, 1083–1094 (2015).

45. Brown DR. Nitrogen Starvation Induces Persister Cell Formation in Escherichia coli. J Bacteriol 201, (2019).

46. Hansen S, Lewis K, Vulic M. Role of global regulators and nucleotide metabolism in antibiotic tolerance in Escherichia coli. Antimicrob Agents Chemother 52, 2718–2726 (2008).

47. Mok WW, Orman MA, Brynildsen MP. Impacts of global transcriptional regulators on persister metabolism. Antimicrob Agents Chemother 59, 2713–2719 (2015).

48. Wu Y, Vulic M, Keren I, Lewis K. Role of oxidative stress in persister tolerance. Antimicrob Agents Chemother 56, 4922–4926 (2012).

49. Xu T, Wang XY, Cui P, Zhang YM, Zhang WH, Zhang Y. The Agr Quorum Sensing System Represses Persister Formation through Regulation of Phenol Soluble Modulins in Staphylococcus aureus. Front Microbiol 8, 2189 (2017).

50. Cui P, Niu H, Shi W, Zhang S, Zhang W, Zhang Y. Identification of Genes Involved in Bacteriostatic Antibiotic-Induced Persister Formation. Front Microbiol 9, 413 (2018).

51. García-Del Portillo F, Pucciarelli MG, Casadesús J. DNA adenine methylase mutants of Salmonella typhimurium show defects in protein secretion, cell invasion, and M cell cytotoxicity. Proc Natl Acad Sci USA 96, 11578–11583 (1999).

52. Low DA, Casadesús J. Clocks and switches: bacterial gene regulation by DNA adenine methylation. Curr Opin Microbiol 11, 106–112 (2008).

53. Nicholson B, Low D. DNA methylation-dependent regulation of pef expression in Salmonella typhimurium. Mol Microbiol 35, 728–742 (2000).

54. Adhikari S, Curtis PD. DNA methyltransferases and epigenetic regulation in bacteria. FEMS Microbiol Rev 40, 575–591 (2016).

55. Kahramanoglou C, et al. Genomics of DNA cytosine methylation in Escherichia coli reveals its role in stationary phase transcription. Nat Commun 3, 886 (2012).

56. Donczew R, Zakrzewska-Czerwińska J, Zawilak-Pawlik A. Beyond DnaA: the role of DNA topology and DNA methylation in bacterial replication initiation. J Mol Biol 426, 2269–2282 (2014).

57. Mok WWK, Brynildsen MP. Timing of DNA damage responses impacts persistence to fluoroquinolones. Proc Natl Acad Sci U S A 115, E6301–E6309 (2018).

58. Lv H, Li J, Wu Y, Garyali S, Wang Y. Transporter and its engineering for secondary metabolites. Appl Microbiol Biotechnol 100, 6119–6130 (2016).

